# State-dependent alteration of respiration in a rat model of Parkinson disease

**DOI:** 10.1101/2023.11.10.566528

**Authors:** Jean Jacques Walker, Estelle Meunier, Samuel Garcia, Belkacem Messaoudi, Anne-Marie Mouly, Alexandra Veyrac, Nathalie Buonviso, Emmanuelle Courtiol

## Abstract

Parkinson disease (PD) is the second most frequent neurodegenerative disorder. Besides major deficits in motor coordination, patients may also display sensory and cognitive impairments, which are often overlooked despite being inherently part of the PD symptomatology. Amongst those symptoms, respiration, a key mechanism involved in the regulation of multiple physiological and neuronal processes, appears to be altered. Importantly, breathing patterns are highly correlated with the animal’s behavioral states, and although respiration has been investigated in different models of PD, no study has yet taken into consideration the potential impact of behavioral state on respiration deficits in these models. To explore this variable, we first characterized the respiratory parameters in a neurotoxin-induced rat model of PD (6-OHDA) across different vigilance states: sleep, quiet waking and exploration. We noted a significantly higher respiratory frequency in 6-OHDA rats during quiet waking compared to Sham rats. A higher respiratory amplitude was also observed in 6-OHDA rats during both quiet waking and exploration. No effect of the treatment was noted during sleep. Given the relation between respiration and olfaction and the presence of olfactory deficits in PD patients, we then investigated the odor-evoked sniffing response in PD rats, using an odor habituation/cross-habituation paradigm. No substantial differences were observed in olfactory abilities between the two groups, as assessed through sniffing frequency. These results corroborate the hypothesis that respiratory impairments in 6-OHDA rats are vigilance-dependent. Our results also shed light on the importance of considering the behavioral state as an impacting factor when analyzing respiration.

**Highlights:**

- Specific respiratory patterns associated to different vigilance states
- Specific alteration of respiration during quiet waking in a rodent model of PD
- Preserved olfactory abilities as assessed through sniffing in a rodent model of PD

## Introduction

Parkinson’s disease (PD) is an insidious and progressive neurodegenerative disorder. It is characterized by an extensive loss of dopaminergic neurons in the substantia nigra pars compacta [SNc; (Poewe et al. 2017)]. This structure is known to send ascending dopaminergic outputs to subcortical targets. Their subsequent degeneration notably leads to a massive loss of the striatal outputs, therefore resulting in well-characterized movement disorders such as rest tremor, bradykinesia, rigidity and postural instability. In addition, this pathology is also believed to affect other brain areas such as the locus coeruleus, the olfactory system, the limbic cortex and the peripheral nervous system (Braak et al. 2003; Braak et al. 2007; Mu et al. 2013). Beside the motor features, it is therefore common that PD patients display non-motor symptoms, including pain, sleep disorders, cognitive and emotional impairments as well as olfactory deficits (Castrioto et al. 2016; Doty et al. 1988; Drui et al. 2014; Poewe 2008; Roos et al. 2019; Schapira et al. 2017; Sobel et al. 2001; Sung and Nicholas 2013), which often occur prior to the onset of the motor symptoms. Not only do they contribute to the decline of the patient’s health-related quality of life, but also several reports have suggested that they could be more debilitating than the motor symptoms (Breen and Drutyte 2013; Hinnell et al. 2012). Yet, those non-motor symptoms have remained neglected for a long time in PD patients’ care considering their unspecific symptomatology.

Amongst those symptoms, issues associated with the respiratory functions are reported in PD patients. Respiration is a vital function that, in addition to supplying oxygen to the body, regulates multiple physiological and neural processes. Due to its rhythmical nature, respiration has been recently demonstrated to entrain brain neural oscillatory activity at its frequency and to synchronize high frequency oscillatory activities of various brain areas such as the olfactory areas, the hippocampus or the prefrontal cortex [for review: (Heck et al. 2019; Juventin et al. 2023; Kluger and Gross 2021; Tort et al. 2018)]. Those different oscillatory rhythms are crucial for perception, adaptive motor responses, and memory formation (Fries 2005, 2015; Tallon-Baudry et al. 1999). Accordingly, respiration has been shown to impact detection of fear expressing faces as well as performance in a visuo-spatial task (Perl et al. 2019; Zelano et al. 2016) but also consolidation of episodic odor memory (Arshamian et al. 2018). PD-focused studies have reported various respiratory symptoms in patients such as upper airway obstruction, tachypnea, dyspnea or sleep apnea [(Apps et al. 1985; Baille et al. 2016; Docu Axelerad et al. 2021; Torsney and Forsyth 2017); see reviews: (Aquino et al. 2022; Kaczyńska et al. 2022)]. In order to better understand those respiratory deficits, their temporal decay as well as their underlying mechanisms, different studies have used rodent models of PD and recorded their respiration under normoxic, hypoxic and hypercapnic conditions. The effects reported were quite variable depending on the study (Aquino et al. 2022; Kaczyńska et al. 2022). For instance, under normoxic conditions, some studies highlighted a decrease in respiratory frequency in rat models of PD compared to controls (Andrzejewski et al. 2020; Bialkowska et al. 2016; Falquetto et al. 2020; Fernandes-Junior et al. 2018; Lima et al. 2018; Oliveira et al. 2017, 2019; Tuppy et al. 2015) while another study showed an increase in respiratory frequency (Johnson et al. 2020b). Interestingly, other studies did not observe any difference (Andrzejewski et al. 2016, 2019). Under hypoxic conditions, some studies demonstrated a decrease in respiratory frequency in rats’ models of PD (Andrzejewski et al. 2016; Fernandes-Junior et al. 2018) while others did not observe any difference (Johnson et al. 2020b; Oliveira et al. 2017; Tuppy et al. 2015). For hypercapnic conditions, the results were more homogenous and showed a decreased respiratory frequency in rats’ model of PD compared to Sham rats (Andrzejewski et al. 2016, 2019; Fernandes-Junior et al. 2018; Johnson et al. 2020b; Lima et al. 2018; Oliveira et al. 2017; Tuppy et al. 2015). Those different respiratory effects seem to originate in a loss of neurons in the respiratory centers (Falquetto et al. 2017, 2020; Fernandes-Junior et al. 2018; Lima et al. 2018; Oliveira et al. 2019, 2021; Tuppy et al. 2015). Kaczyńska et al. (2022) proposed that the inconsistency in the results obtained on respiration in rat models of PD is due to the use of different models of PD and experimental conditions. Importantly, none of these studies seem to have taken into consideration the animal behavioral state while recording their respiration. Of note, different teams have shown that different behaviors are associated with specific respiratory patterns (Bagur et al. 2018, 2021; Dupin et al. 2020; Girin et al. 2021; Hegoburu et al. 2011; Janke et al. 2022; Youngentob et al. 1987). For instance, major differences in respiratory frequency and peak flow rate can be observed between sleep, quiet waking or exploration (Girin et al. 2021). The effect on respiration in rat models of PD might thus also be related to different vigilance states expressed. Alternatively, respiratory alterations observed in PD rat models may also be specific of a given behavioral state. To disentangle these points, we first investigated the behavioral states expressed in a neurotoxin-induced rat model of PD and tested whether respiratory alterations observed in rat model of PD could be specific of a given vigilance state (sleep, quiet waking and exploration).

Respiration acts as a mechanical vector of odorant molecules. Active sensory sampling is particularly prominent in rats and has been shown to influence the olfactory system neural activity and functions (Buonviso et al. 2006; Juventin et al. 2023; Mainland and Sobel 2006). From a clinical standpoint, it is important to note that 90% of PD patients display moderate to severe olfactory impairments, which can occur as early as the prodromal stage of the disease (Doty et al. 1988; Fullard et al. 2017; Haehner et al. 2011). Interestingly, PD patients suffering from disrupted smell identification and odor threshold detection tend to also display impaired sniffing abilities (Sobel et al. 2001). Given the inextricable relation between respiration and olfaction, the second part of our study aimed at characterizing the olfactory abilities of our PD rats. We explored the rats’ odor-evoked sniffing response in an odor habituation/cross-habituation paradigm, a perceptual and mnesic task testing olfactory abilities.

A neurotoxin-induced model of PD consisting of inducing a selective and bilateral lesion of the SNc using 6-hydroxydopamine (6-OHDA) was used to conduct our experiments. As described by Carnicella et al. (2014), this model is not only convenient for reproducing non-motor symptoms of the disease but also for circumventing the motor disorders. A whole-body plethysmograph was used to record the respiration of 6-OHDA and Sham rats in a range of 3-5 weeks following induction of the lesion. We first noted that the amount of sleep, exploration and quiet waking was not different between 6-OHDA and Sham rats. Then, we demonstrated that respiratory parameters were not impacted by the 6-OHDA treatment during sleep. However, we observed significant effects of the treatment on respiration during the awake states: 6-OHDA rats displayed higher respiratory frequency than Sham rats during quiet waking and higher respiration amplitude during both quiet waking and exploration. During the odor habituation/cross-habituation task, we did not observe any significant difference in sniffing frequency nor olfactory abilities between the two groups. These results highlight the specific alteration of 6-OHDA rats’ respiration notably during the awake states. Our results also unveil the critical importance of considering the behavioral state while recording respiration.

## Material and Methods

### Animals

Adult male Sprague-Dawley rats (n = 28, Charles River, L’Arbresle, France) weighing between 200 and 250 grams at their arrival in the animal facility (6 weeks old) were used. Animals were housed in pairs, under controlled optimum conditions including temperature 22°C ± 1 °C, hygrometry 50% ± 10%, constant light / dark cycle 7 am to 7 pm with food and water *ad libitum*. The experiments were carried out according to the ethical guidelines of the European Communities Council Directive 2010/63/EU, as well as the approval #16979 of the Lyon 1 University CEEA-55 ethical committee and of the Ministry of Higher Education, Research and Innovation. All efforts were made to minimize the number of animals used and their suffering. Each experiment was designed in agreement with the “3R” rules.

### Bilaterally lesioned 6-OHDA rat model of Parkinson’s Disease

In order to facilitate manipulations and to reduce handling-related stress, rats were habituated to human contact prior to the experiments. For a week, all rats were handled and familiarized to the experimenter’s scent and voice, for a minimum of 15 minutes per day.

### Surgical procedure

Surgical interventions were performed 4 weeks after the rats’ arrival into the animal facility. Animals were assigned to either the 6-OHDA (n=16) or the Sham group (n = 12), ensuring that the weight average was similar across both groups before the surgery. Rats were anesthetized with isoflurane gas using an induction chamber (4%, combined with O_2_, Compact anesthesia module, Minerve, Esternay France) and then placed in a stereotaxic apparatus under constant isoflurane anesthesia on a heating pad. The animal received an injection of Carprofen (5mg/kg, s.c., ZOETIS France SAS) for analgesia. In addition, an injection of desipramine (s.c., 15 mg/kg, Sigma Aldrich) was made 30 minutes before the intracerebral injections in order to protect the noradrenergic and serotoninergic neurons from the neurotoxic effect of 6-OHDA, as well as to increase its selectivity and efficacy (Carnicella et al. 2014; Drui et al. 2014). Two holes were drilled in the skull over the SNc according to the following stereotaxic coordinates relative to the Bregma : AP : - 5.4 mm; ML : ± 1.8 mm; DV : - 8.1 mm.

Using a 10 μL Hamilton syringe coupled with a microneedle-silica fiber (75µm internal diameter and 150µm external diameter, Phymep) set on a micro-syringe pump injector (UMP3 UltraMicroPump, WPI), 6-OHDA rats received 2.3 μL bilateral injections of 6-OHDA hydrobromide (6 μg combined with ascorbic acid dissolved in 2.3 μL sterile 0.9% NaCl, Sigma Aldrich) in the SNc in each cerebral hemisphere, whereas Sham rats receive 2.3 μL saline (NaCl, 0.9%) at 0.5 μL/min rate. Upon completion of administration of 6-OHDA or saline, the needle was left in place for a further ten minutes to allow diffusion of the solution in the area. After removing the needle, a bio-compatible, non-toxic and rapid curing silicone adhesive (Kwik-Cast™) was used to seal the two bone holes. Stitches were made using antibacterial Vicryl*Plus suture (Ethicon®).

After recovering from anesthesia, rats were returned to the animal facility and were monitored daily for at least 3 weeks, allowing the dopaminergic lesion to extend. Their weight, posture, coat condition, motor skills and general state were examined at least once a day. Following surgery, rats were supplied with highly caloric and palatable food and received appropriate nursing. One of the 6-OHDA rats did not recover from the surgery leaving a total n of 15 6-OHDA and 12 Sham rats.

### Respiration and behavior recordings

#### Whole-body plethysmograph

Respiration was measured by using a whole-body plethysmograph [diameter: 20cm, height: 30cm, EMKA Technologies, France ; (Courtiol et al. 2014; Lefèvre et al. 2016)]. It consists in a plexiglass cage composed of two independent chambers: the recording one where the rat is installed and the reference one. The changes in pressure resulting from the animals’ breathing were detected using a differential pressure transducer located between the recording chamber and the reference one. The signal was sampled at 1 kHz, amplified (Amplipower, EMKA Technologies) and recorded using a home-made software developed in Python (pyacq, https://github.com/pyacq/pyacq). A constant airflow was delivered from the top (1100 ml/min) and from a central port situated on the wall of the plethysmograph (400ml/min) for a total of 1500 mL/min. The air was vacuumed out at the bottom at 1500 mL/min to ensure constant ventilation. During the odor-evoked paradigm, pre-selected pairs of odors were presented via an olfactometer through the central port. The whole-body plethysmograph was placed in an artificially enlighten sound-attenuating enclosure, isolating the rat from the surrounding environment. The entire experiment was video recorded via a webcam (HD Webcam C930e) mounted on the side wall of the sound-attenuating enclosure. Videos were captured using the pyacq software and synchronized to the respiratory signal for each session.

#### Experimental Paradigms

Animals were first gradually habituated to the experimental cage for a week prior to starting any recording. A week before surgery, rats underwent recording sessions (Pre-Surgery; Fig. 1) to establish a baseline. Post-Surgery recordings then started three to four weeks following the surgery, and occurred on a weekly basis for two consecutive weeks (S1 and S2; Fig. 1). During each of those recording weeks, each rat underwent two sessions: the first with a passive exposure to the experimental cage to test for the effect of the vigilance state and the second one with odors in the habituation/cross-habituation paradigm (Fig. 1).

**Figure 1:**
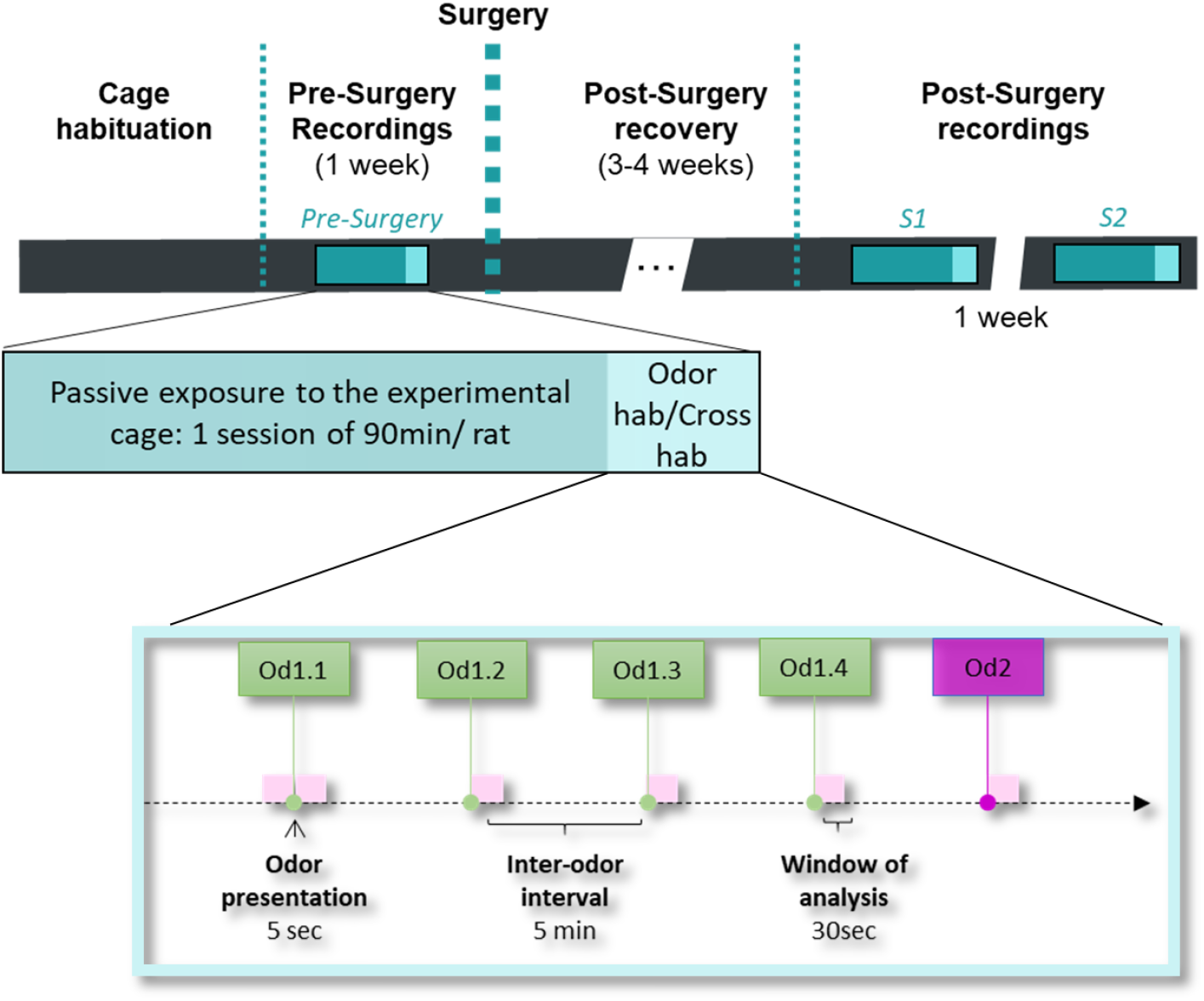
Timeline of the experimental study. Rats were first habituated to the cage, then Pre-Surgery recordings were performed: one day for a passive exposure to the experimental cage session (green) and one day for the odor protocol (light green; Od1 is presented four times followed by the presentation of Od2). The rats then underwent surgeries and were allowed to recover for at least three weeks before the Post-Surgery recordings. As for the Pre-Surgery recording week, each rat underwent two sessions: one day for a passive exposure to the experimental cage and one day for the odor habituation/cross-habituation task .

##### Passive exposure to the experimental cage

Once a week, each rat was introduced in the whole-body plethysmograph and allowed a 90-minute period of free exploration. Different behaviors, such as grooming, rearing, resting and sleeping were observed. There was no task to perform, nor odor diffused.

##### Odor-evoked sniffing: odor habituation/cross-habituation paradigm

Once a week, each rat was placed in the experimental chamber for 30 minutes and odors were presented in a habituation / cross-habituation paradigm (Al Koborssy et al. 2019). This paradigm is a perceptual and mnesic task allowing to study odor detection, habituation, discrimination, and in which sniffing frequency has been shown to be a useful and sensitive readout of olfactory abilities (Al Koborssy et al. 2019; Coronas-Samano et al. 2016; Johnson et al. 2020a).

After five minutes in the plethysmograph allowing the rat to get to a more relaxed state, a first odor was presented four times (Od1.1, Od1.2, Od1.3 and Od1.4), followed by the presentation of a second odor (Od 2). Each odor was presented during 5 seconds at 400 mL/min thanks to a semi-automatic setup, with an inter-odor interval of 5 minutes (Fig. 1; bottom). For our study, all odorants used were unfamiliar and at saturated vapor pressure. The pairs of odors were different each week with ethyl-heptanoate (Od1) / ethyl-hexanoate (Od2) for the Pre-Surgery session, carvone (Od1) / Citral (Od2) for S1 session and limonene (Od1) / anethol (Od2) for S2 session (Sigma-Aldrich).

### Data analysis

#### Respiratory signal

Respiration appears to be a periodic function associating negative (inspiration) and positive (expiration) deflections. The analysis of the respiratory data relied on a python-coded algorithm allowing the detection of the transition point between inspiration and expiration (Courtiol et al. 2014; Girin et al. 2021; Lefèvre et al. 2016; Roux et al. 2006). Inspiration was considered to start at the zero-crossing point of the falling phase and ends at the zero-crossing point of the rising phase, whereas the expiratory phase began at the zero-crossing point of the rising phase and ends at the zero-crossing point of the falling phase.

In order to reduce noise and to eliminate potential artifacts, raw data were visually inspected, and an automatic Python script was further used to remove artifacts from the dataset. Cut-off values were determined upon the 5 first minutes of each recording. The median and median absolute deviation of the inspiration amplitude were estimated for each rat and session over that 5-minute period. Each cycle for which value was above the threshold of median + 8x median absolute deviation was discarded. This threshold value was based on our observations and allowed to dissociate respiratory cycles from artifacts. When artifacts were separated by less than a 10-second period, neighboring cycles were agglomerated into batches which were also discarded. Following the detection, values of respiratory cycle parameters were extracted. We notably analyzed the instantaneous frequency (1/cycle duration; Hz) and the total amplitude (representing the sum of the maximal inspiratory and expiratory peak flow rates) normalized to the animal’s weight (ml/sec/100g).

##### Respiratory signals in different behavioral states

The association between animal behavior and their specific respiratory outcomes has been described in several studies (Dupin et al. 2020; Girin et al. 2021; Hegoburu et al. 2011; Janke et al. 2022). Therefore, during the passive exposure to the experimental cage, we decided to inspect the rats’ respiratory signal across three vigilance states: sleep, quiet waking and exploration (Girin et al. 2021). The video recordings of each session were manually encoded using the ephyviewer (https://github.com/NeuralEnsemble/ephyviewer.git). Sleeping was encoded when the rat had his eyes closed, with his body lying on the cage floor or in a curled-up position (Fig. 2B). The quiet waking state was encoded when the rat’s eyes were open and when he was either immobile or slightly moving. Exploration was defined when the rat was clearly exploring its environment or standing on his two back feet (rearing). Behaviors such as grooming, state transition or undefined activities (e.g. when the rat’s eyes were not clearly visible) were not encoded and discarded from the analysis. The mean value of the different respiratory parameters was measured per animal and session under each behavioral state.

**Figure 2:**
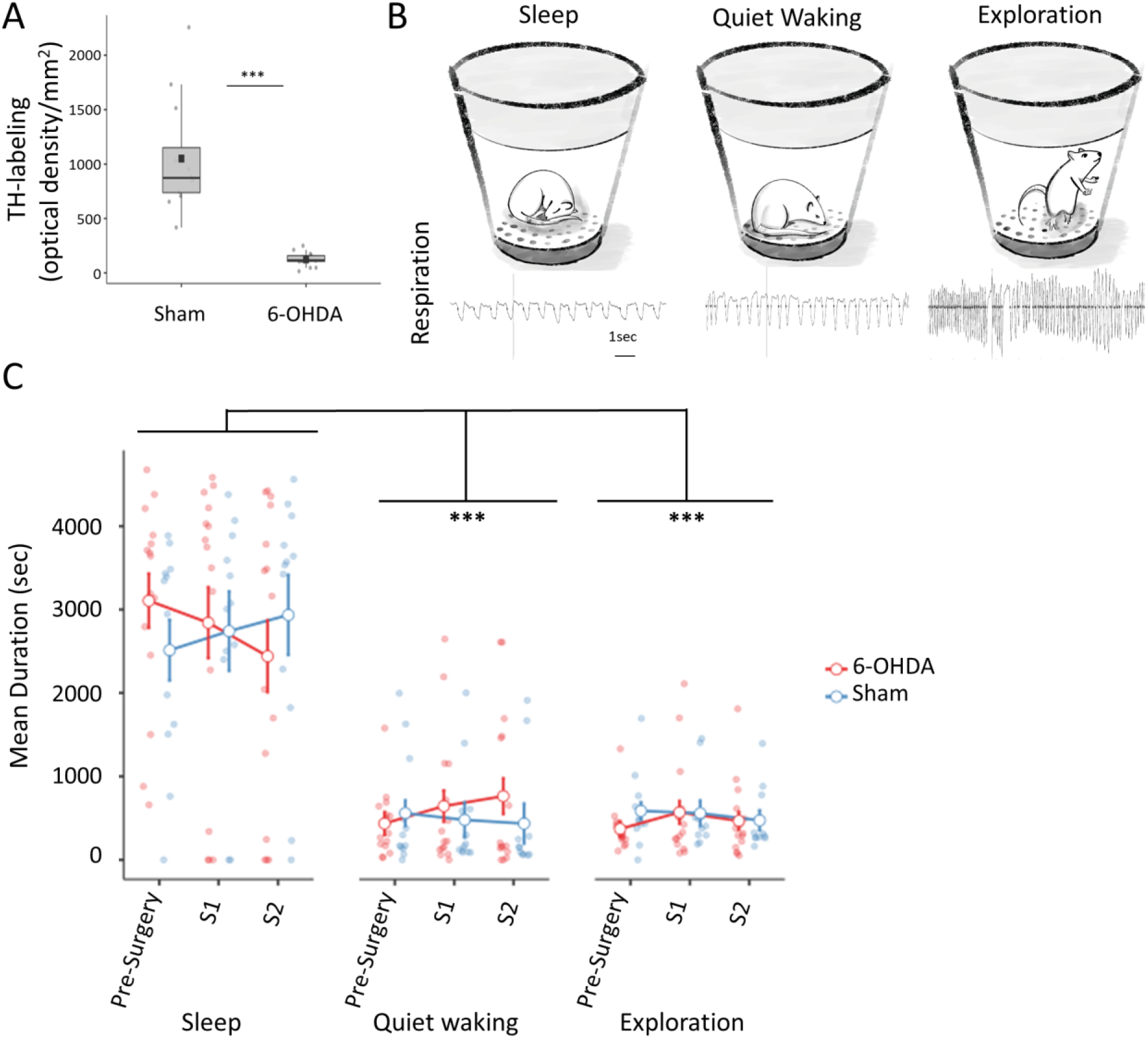
A-Box plot of the quantification of the loss of TH-labeling through optical density/mm^2^ in 6-OHDA rats compared to Sham rats. Unpaired t-test, *** p<0.001. n 6-OHDA = 13 and n= 12 Sham rats. Black squares represent the mean value of each group. B-Examples of the three vigilance states (top) and their associated respiratory pattern (below) with from left to right: sleep, quiet waking and exploration. There are clear differences in terms of respiratory frequency and total amplitude. C-Mean (±sem) duration of the time spent during each state (sleep, quiet waking and exploration) for each session (Pre-Surgery, S1 and S2) and for 6-OHDA rats (n=15; red) and Sham rats (n=12; blue). Repeated Measures ANOVA and Fisher’s PLSD post-hoc comparison tests, when significant *** p<0.001.

##### Odor-evoked sniffing response in the odor habituation/cross-habituation paradigm

Using the odor habituation / cross-habituation paradigm, we assessed 1) the odor-evoked sniffing in response to the first presentation of an odor (odor detection), 2) how this response evolves over repetition of that same odor (odor habituation and short-term memory processes) and 3) the sniffing response to a new odor (odor discrimination; (Al Koborssy et al. 2019)). Respiratory signals were analyzed over a 30-second window prior to the first presentation of odor 1 (Pre-Od1; Fig. 1 bottom) and following each odor presentation (30s.; Od1.1; Od1.2; Od1.3; Od1.4 and Od2). Mean values of instantaneous respiratory frequency (Hz) were calculated per animal and per period over this 30s-time window.

### Histology and immunohistochemistry analysis

After the experiments, animals were anesthetized with isoflurane and injected with a lethal dose of Euthasol (400mg/kg). Animals were then transcardially perfused with 0.9% saline and 4% paraformaldehyde. Brains were removed and stored in a paraformaldehyde (4%) solution overnight. Brains were then placed in a 30% sucrose solution for 2 and a half days, frozen (−35°C/40°C) and stored at −80°C. Coronal serial sections of the brains from the midbrain (20µm thick and spaced by 60µm) were performed using a cryostat (Leica CM1950). Those sections were processed for tyrosine hydroxylase (TH) immunohistochemistry in order to label dopaminergic neurons. Brain sections were first rinsed with PBS then treated with 0.5% Triton in PBS for 30 min. Endogenous peroxidases were blocked using a 3% H_2_O_2_ solution in 0.1 M PBS. Sections were then incubated for 90 minutes in 5% normal horse serum (Ozyme), 1% BSA and 0.1% Triton to block nonspecific binding. Sections were incubated overnight at room temperature in a primary rabbit tyrosine hydroxylase antibody (1/300; P40101, Pel-Freez). After incubation for 120min at room temperature in a horse anti-rabbit secondary antibody (1/200; BA-1100, Vector laboratories), sections were processed with avidin-biotin-peroxidase complex (1/200 ; Vectastain Elite ABC-HRP kit, Eurobio) for 30 min. Finally, peroxidase detection was conducted with 0.066% of 3,3-diaminobenzidine-tetrahydrochloride (DAB; Sigma-Aldrich), 0.03% NiCl_2_ and 0.06% H_2_O_2_. Sections were dehydrated in graded ethanol and cover-slipped with Depex (Sigma-Aldrich).

Brain sections were examined with an optical microscope (Axio Scope A1, ZEISS) coupled to a stereology station (Explora Nova Mercator Pro). Quantitative imaging analyses of TH-labeling were performed at 40X magnification in semiautomatic mode. We measured dopaminergic neuronal optical densities bilaterally (left and right) on two sections of the SNc (approximately −5.2/5.4mm and −5.8/−6.04mm posterior from bregma). The mean optical density of TH was calculated per mm^2^ and averaged for the two sides and the two sections. The values were averaged for each experimental group (Sham and 6-OHDA). Due to poor brain quality tissue, two 6-OHDA rats were excluded from histology analysis.

### Statistical analysis

All statistical analyses were performed using Statview 5.01. TH-labeling was compared between groups using an unpaired t-test. A three-way repeated measures ANOVA was used to compare the states durations between the two groups (6-OHDA vs Sham; independent measure) and across sessions and vigilance states (Pre-Surgery, S1 and S2 as well as sleep, quiet waking and exploration; dependent measures). The ANOVA was followed by Fisher’s PLSD post-hoc comparisons when significant.

During passive exposure to the experimental cage, there was a strong difference in respiratory pattern associated with each behavioral state. In addition, some animals did not display a specific state (i.e. sleep, quiet waking or exploration) during at least one of the sessions, thus resulting in a different number of animals for each vigilance state. Respiratory parameters were therefore compared independently for each behavioral state by using a two-way repeated measures ANOVA with the group (6-OHDA vs Sham) as an independent factor and the sessions (Pre-Surgery, S1 and S2) as the repeated measures. The ANOVA was followed by Fisher’s PLSD post-hoc comparisons.

In the odor habituation/cross-habituation paradigm, a two-way repeated measures ANOVA with the group (6-OHDA vs Sham) as the independent factor and the period (Pre-Od1, Od1.1, Od1.2, Od1.3, Od1.4 and Od2) as the repeated factor was performed for each session (each session having a different pair of odors). The ANOVA was followed by Fisher’s PLSD post-hoc comparisons when significant. Results are presented as mean ± SEM. P values smaller than 0.05 were considered to be significant. Graphics were performed under Jamovi.

## Results

The aim of this study was to investigate the vigilance states expressed in 6-OHDA rats compared to Sham rats and to determine their respiratory parameters under different behavioral conditions. First, we explored respiratory parameters when the rats were simply exposed to the whole-body plethysmograph (no odor nor task) and exhibiting different vigilance states from sleep to exploration. Second, we recorded odor-evoked sniffing during a perceptual and mnesic task testing olfactory abilities.

### Validation of the lesion of SNc dopaminergic neurons

First, we verified that the 6-OHDA injection effectively induced a significant reduction in TH-labeling in the 6-OHDA group compared to the Sham group. We observed a strong decrease of dopaminergic neuronal optical densities in the SNc of 6-OHDA rats compared to Sham rats (unpaired t-test (t(23)=-6.335, <0.0001), validating this experimental model of PD (Fig. 2A).

### Amount of time spent in each behavioral state

We recorded behavior and spontaneous respiration of unrestrained rats a week before surgery (Pre-Surgery) and during two Post-Surgery sessions (S1 and S2) during a passive exposure to the experimental cage. We first examined whether the 6-OHDA treatment had an effect on the amount of time spent in the different vigilance states : sleep, quiet waking and exploration (Fig. 2B-C). A three-way ANOVA revealed no significant effect of group, session nor group X session interaction (group: F(1,25)=0.326, p=0.5729; session: F(2,50)=0.652, p=0.5253; groupXsession F(2,50)=1.434, p=0.248). However, there was a significant effect of state (F(2,50)=39.493; p<0.0001) and post-hoc comparisons revealed that the amount of time spent in sleep state was significantly higher than quiet waking and exploration states for all sessions and for both groups. Indeed, there was no significant interaction between state and group, state and session or state, session and group (state X group: F(2,50)=0.059, p=0.9428; session X state: F(4,100)=0.195, p=0.9404); session X state X group: F(4,100)=2.329, p=0.0612).

### Respiration of 6-ODHA rats during specific vigilance state

We investigated respiration under three different vigilance states: sleep, quiet waking, and exploration (Fig. 2B). We analyzed two respiratory parameters that globally characterize a respiratory cycle: the instantaneous frequency (Hz) and the total amplitude (ml/sec/100g). Values were averaged per rat per session and per state.

#### Sleep

Two-way (group, session) ANOVA did not reveal any group effect nor session X group interaction effect for the two respiratory parameters tested (see table 1). Only a session effect was observed for the instantaneous respiratory frequency (Table 1; Fig. 3A). Post-hoc comparisons revealed a significant difference between the Pre-Surgery session and the two Post-Surgery sessions with a decrease of instantaneous frequency from on average 1.6Hz to 1.3Hz (no difference between S1 and S2; Fig. 3A). Total amplitude did not significantly change across sessions (Fig. 3B).

**Figure 3:**
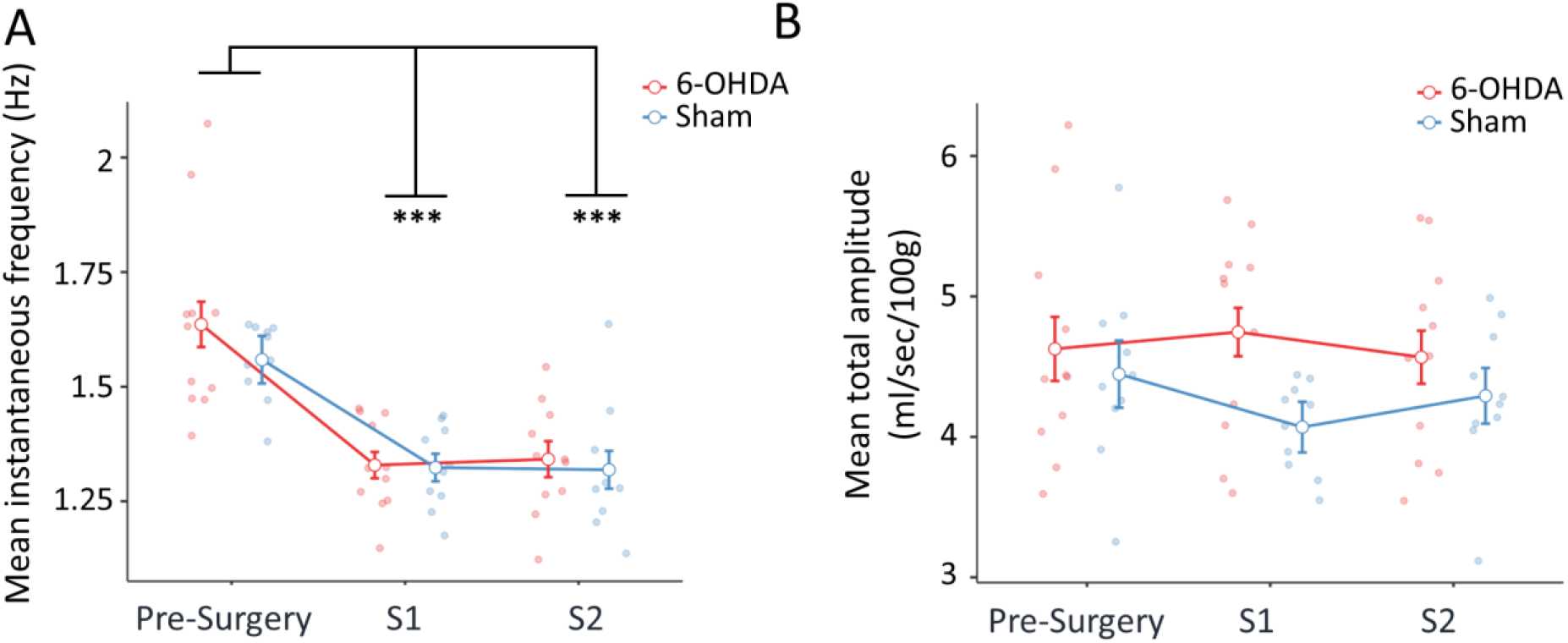
Respiratory parameters were not altered in 6-OHDA rats during the sleep state. A-Mean (±sem) instantaneous respiratory frequency (Hz) and B-Mean (±sem) respiratory total amplitude (ml/sec/100g) across sessions of 6-OHDA rats (n= 11; red) and Sham rats (n= 10; blue). Repeated Measures ANOVA, Post-hoc comparisons with Fisher’s PLSD test when significant; *** p<0.001.

**Table 1:**
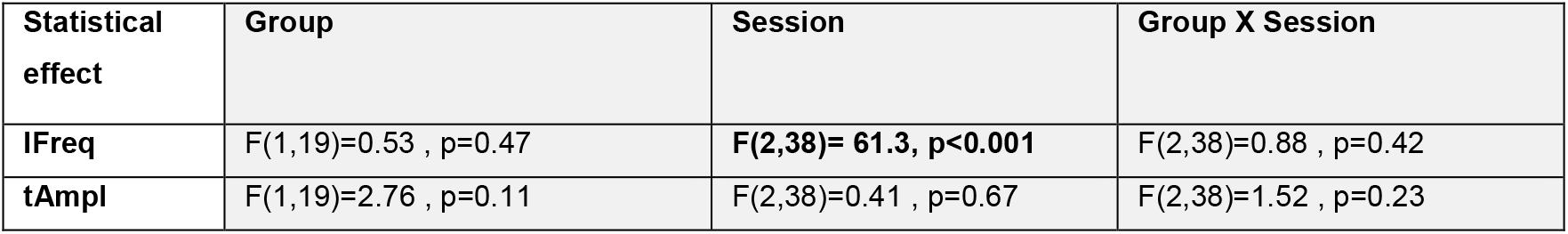
Statistical effects on respiratory parameters during the Sleep state. Repeated Measures ANOVA with the session as a repeated factor and the group as the independent factor. IFreq = instantaneous respiratory frequency (Hz) and tAmpl = total amplitude (ml/sec/100g).

| Statistical effect | Group | Session | Group X Session |
| --- | --- | --- | --- |
| IFreq | $F(1,19)=0.53$ , $p=0.47$ | **$F(2,38)= 61.3$ , $p<0.001$** | $F(2,38)=0.88$ , $p=0.42$ |
| tAmpl | $F(1,19)=2.76$ , $p=0.11$ | $F(2,38)=0.41$ , $p=0.67$ | $F(2,38)=1.52$ , $p=0.23$ |

Altogether, our results show an evolution in respiratory frequency between the Pre-Surgery session and both Post-Surgery sessions with a lowering of respiratory frequency during sleep. This decrease might be attributed to an habituation to the cage and to longer periods of slow wave sleep over time, respiration frequency being slower during slow wave sleep (Girin et al. 2021). This effect was similar between both groups of rats. Therefore, the 6-OHDA treatment does not significantly impact respiration during sleep. We next explored two awake states.

#### Quiet waking

Two-way (group, session) ANOVA did not reveal a significant group effect for instantaneous frequency however a significant session effect was observed (Table 2). In addition, we also observed a significant interaction between group and session for the instantaneous respiratory frequency (Table 2). Post-hoc analyses revealed a significant decrease in instantaneous respiratory frequency between the Pre-Surgery session and both Post-Surgery sessions in the Sham group, as observed during the sleep state, from 2.4 Hz to 2Hz and 1.9 Hz, while there was no change for the 6-OHDA group (on average 2.3, 2.3 and 2.2 Hz for Pre-Surgery, S1 and S2, respectively; Fig. 4A). As a consequence, when considering each session, post-hoc tests revealed a significant difference between 6-OHDA and Sham rats during S1. While not significant, the same tendency was observed for S2, with a slower instantaneous respiratory frequency in Sham than in 6-OHDA rats.

**Figure 4:**
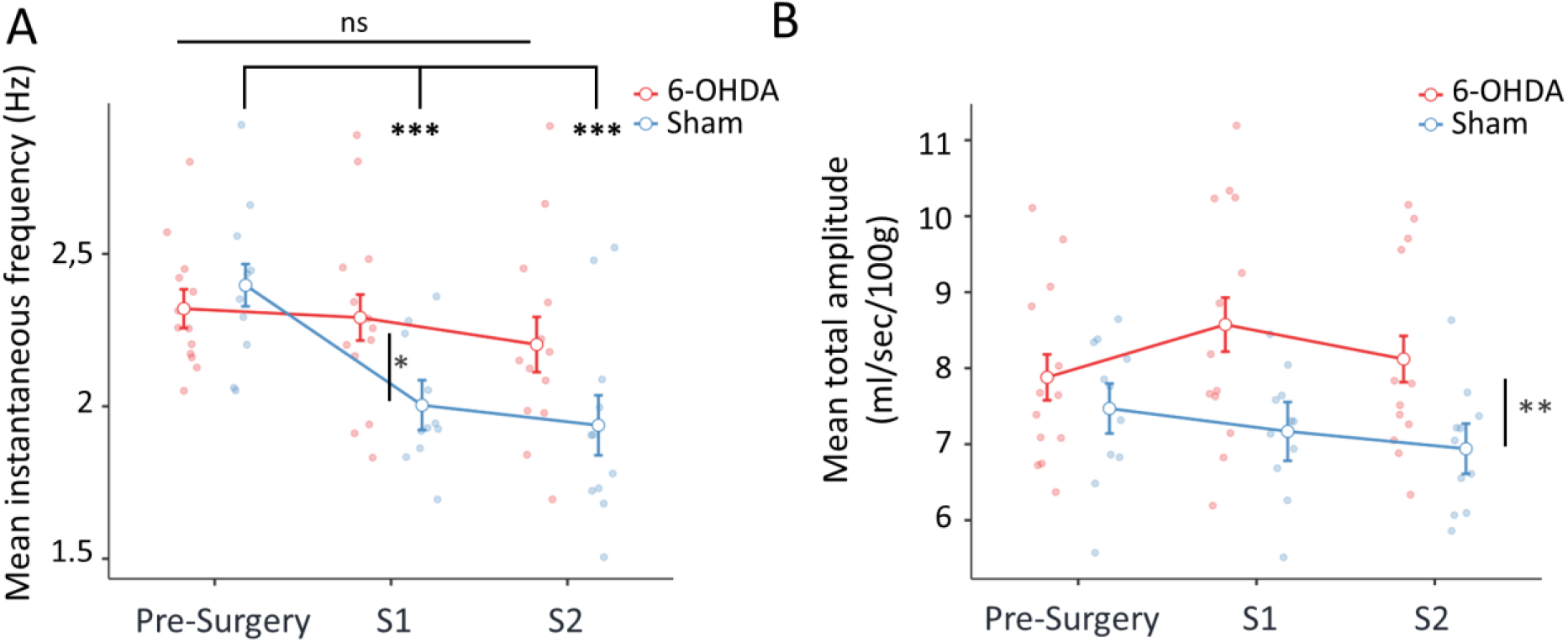
Alteration of respiratory frequency and total amplitude of 6-OHDA rats during the quiet waking state. A-Mean (±sem) instantaneous respiratory frequency (Hz) and B-Mean (±sem) respiratory total amplitude (ml/sec/100g) across sessions of 6-OHDA rats (n= 13; red) and Sham rats (n= 11; blue). Repeated Measures ANOVA, Post-hoc comparisons with Fisher’s PLSD test with a significant difference (*** p<0.001) between the Pre-Surgery session and the Post-Surgery sessions for the Sham group and a significant difference between 6-OHDA and Sham rats during S1 (* p<0.05) for the instantaneous frequency. For the total amplitude, there was a global effect of the group (** p<0.01). .

**Table 2:** Statistical effects on respiratory parameters during the Quiet Waking state. Repeated Measures ANOVA with the session as a repeated factor and the group as the independent factor.

| Statistical effect | Group | Session | Group X Session |
| --- | --- | --- | --- |
| IFreq | $F(1,22)=3.55, p=0.073$ | $F(2,44)=9.93, p=0.0003$ | $F(2,44)=4.64, p=0.015$ |
| tAmpl | $F(1,22)=9.17, p=0.006$ | $F(2,44)=0.67, p=0.52$ | $F(2,44)=1.56, p=0.22$ |

For total amplitude, the Two-way (group, session) ANOVA revealed a significant effect of the group and did not reveal a difference across sessions or group X session interaction (Fig. 4B). 6-OHDA rats displayed a higher respiratory total amplitude compared to Sham rats no matter the session but the figure clearly shows that the effect mainly occurred during the Post-Surgery sessions (as highlighted by the fact that when looking at each session independently, post-hoc comparisons were significant at S1 and S2 but not at Pre-Surgery).

In conclusion, we observed a significant effect of the lesion of the dopaminergic neurons in the SNc on the respiratory frequency and total amplitude during the quiet waking state. 6-OHDA rats notably display higher respiratory frequency and total amplitude compared to Sham rats. We then analyzed another awake state : exploration.

#### Exploration

Two-way (group, session) ANOVA did not reveal any effect on respiratory frequency during the exploration state (see Table 3). The mean instantaneous frequency of sniffing was stable across sessions and between groups with an average respiratory frequency of about 7Hz (Fig. 5A). For the respiratory total amplitude, the repeated measures ANOVA revealed a group and a session X group effect (Table 3). Post-hoc analyses revealed a significant difference between the two groups during S1 with 6-OHDA rats displaying a higher total amplitude than Sham rats, similarly to what was observed during quiet waking. While not significant, a similar tendency was observed at S2. As a result, the two groups also evolved in a different way across sessions, with Sham rats displaying no change of total amplitude across sessions while 6-OHDA rats increasing their total amplitude between the Pre-Surgery session and S1 (Figure 5B).

**Figure 5:**
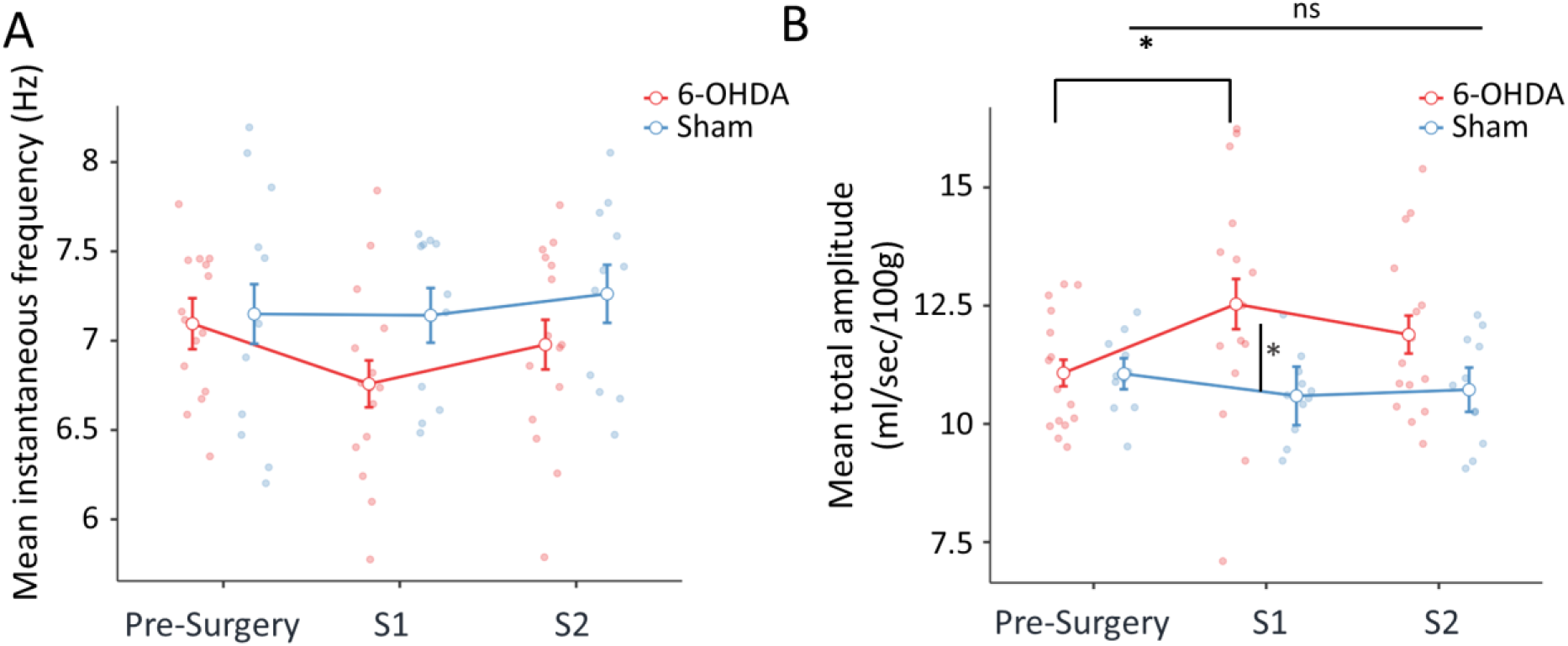
Respiratory parameters were partially altered in 6-OHDA rats during Exploration. A-Mean (±sem) instantaneous respiratory frequency (Hz) and B-Mean (±sem) respiratory total amplitude (ml/sec/100g) across sessions for 6-OHDA rats (n= 15; red) and Sham rats (n= 11; blue). Repeated Measures ANOVA, Post-hoc comparisons with Fisher’s PLSD test with a significant difference (* p<0.05) between the Pre-Surgery session and S1 session for the 6-OHDA group and a significant difference between 6-OHDA and Sham rats during S1 (* p<0.05).

**Table 3:** Statistical effects on respiratory parameters during the Exploration state. Repeated Measures ANOVA with the session as a repeated factor and the group as the independent factor.

| Statistical effect | Group | Session | Group X Session |
| --- | --- | --- | --- |
| IFreq | $F(1,24)=2.07, p=0.16$ | $F(2,48)=1.55, p=0.22$ | $F(2,48)=1.13, p=0.33$ |
| tAmpl | **$F(1,24)=4.75, p=0.04$** | $F(2,48)=0.9, p=0.41$ | **$F(2,48)=3.44, p=0.04$** |

In summary, the respiratory parameters were not affected during sleep but significantly altered during quiet waking. During exploration, our results show that while there was a stability of sniffing frequency across sessions and groups, which might mean that a plateau of sniffing frequency was reached, the total amplitude was significantly impacted by the 6-OHDA treatment.

Importantly, sniffing is part of an orienting response and is triggered whenever the animal explores its environment. However, sniffing during spontaneous exploration as measured previously and sniffing triggered by an odor may not only be different in terms of frequency but the latter also represents an active sensory sampling, which may be specifically altered by the lesion of the dopaminergic system. In the second experiment, we recorded odor-evoked sniffing using a habituation/cross-habituation paradigm, a perceptual and mnesic task that does not require voluntary action from the rats in the whole-body plethysmograph.

### Odor-evoked sniffing during a habituation/cross-habituation paradigm

As shown by others (Al Koborssy et al. 2019; Coronas-Samano et al. 2016) and illustrated in an example in Figure 6A, the first presentation of a novel odor (Od1) induces an increase in sniffing frequency compared to the pre-odor period, reflecting detection of the odor by the animal. This sniffing response decreases across repeated presentation of odor 1, reflecting habituation to the odor as well as short-term memory processes. Finally, the presentation of a second novel odor (Od2) re-initiates a long bout of high-frequency sniffing indicating that the animal discriminates and remembers of the two odors. We compared the mean instantaneous sniffing frequency measured during a 30-sec time window prior to the first odor presentation and after each odor presentation, for both Sham and 6-OHDA rats.

**Figure 6:**
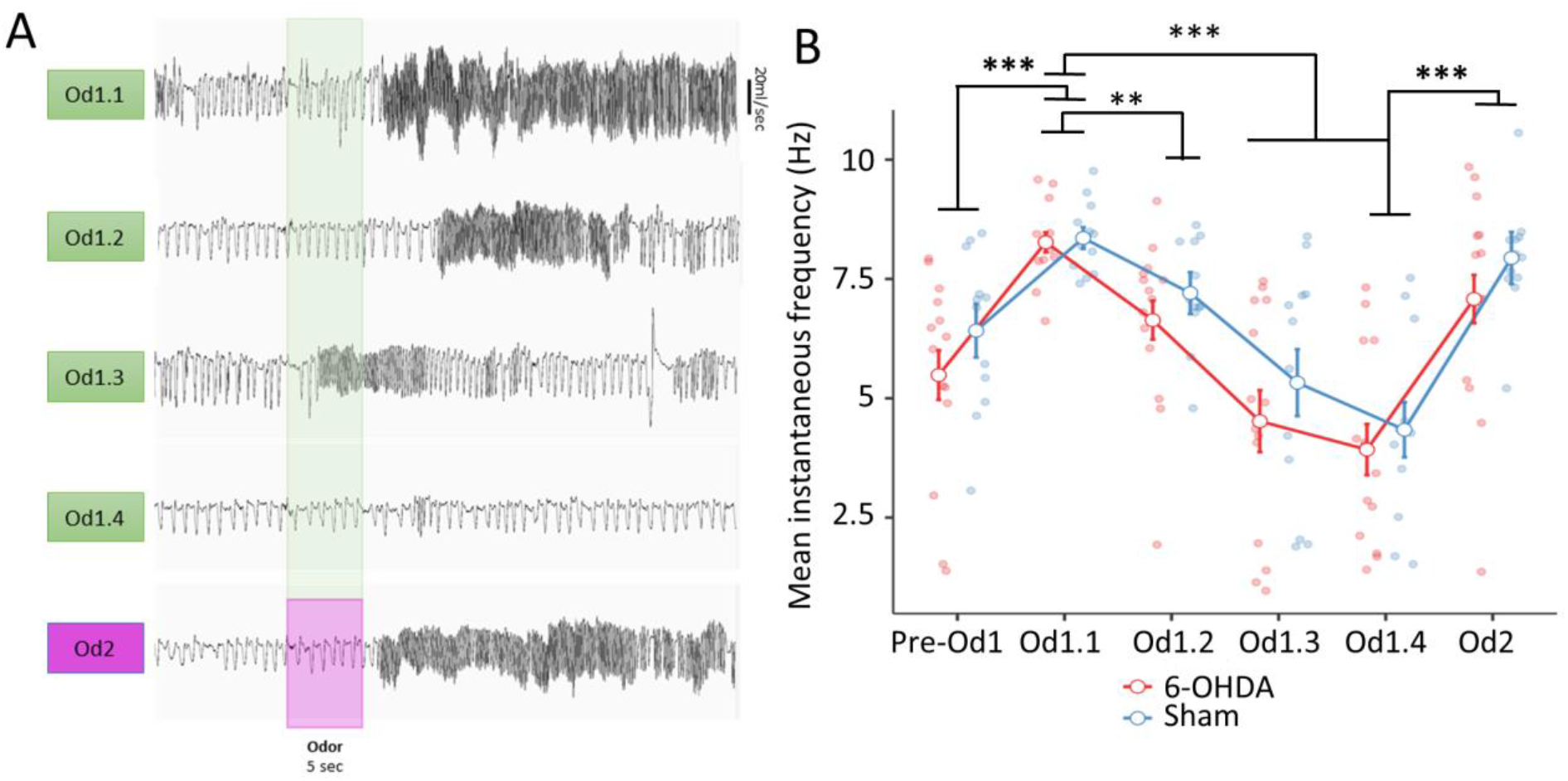
Similar odor-evoked sniffing frequency between 6-OHDA and Sham rats during an odor habituation/cross-habituation task. A-Examples of raw respiratory signals illustrating the respiration baseline followed by the odor-evoked sniffing for each odor. The green cue corresponds to odor 1 presentations (5s; Od1) while the pink cue represents odor 2 (Od2). B-Mean (±sem) instantaneous respiratory frequency (Hz) as a function of the odor period and group. 6-OHDA group: n= 14 (red; there was a technical issue in S1 for one 6-OHDA rat) and Sham group: n = 12 (blue). Values were averaged over 30-sec time bin window for each odor period (before the 1^st^ odor presentation and after the onset of each odor). Repeated Measures ANOVA and post-hoc comparisons with Fisher’s PLSD test with ** p<0.01 and *** p<0.001.

Two-factor repeated ANOVA were carried out for each session separately: Pre-Surgery, S1 and S2. Similar global effects were observed for the three sessions (Table 4). We will thus only detail one of the sessions (S1).

**Table 4:** Statistical effects on respiratory parameters during the odor habituation/cross-habituation paradigm. Repeated Measures ANOVA with the odor periods as a repeated factor and the group as the independent factor. An ANOVA and post-hoc comparisons with Fisher’s PLSD test were performed for each session.*** p<0.001.

| Statistical effect (Instantaneous Frequency, Hz) | Group | Period | Group X Period |
| --- | --- | --- | --- |
| Pre-Surgery (Ethyl-heptanoate – Ethyl hexanoate) | F(1,25)=1.59, p=0.22 | **F(5,125)=36.01 p<0.0001**<br>*Pre-Od1 vs Od1.1 \*\*\**<br>*Od1.1 vs Od1.4 \*\*\**<br>*Od1.4 vs Od2 \*\*\** | F(5,125)=0.99 , p=0.42 |
| S1 (Carvone – Citral) | F(1,24)=1.83, p=0.19 | **F(5,120)=26.8, p<0.0001**<br>*Pre-Od1 vs Od1.1 \*\*\**<br>*Od1.1 vs Od1.4 \*\*\**<br>*Od1.4 vs Od2 \*\*\** | F(5,120)=0.27 , p=0.93 |
| S2 (Limonene-Anethol) | F(1,24)= 3.12, p= 0.09 | **F(5,120)=28.5 , p<0.0001**<br>*Pre-Od1 vs Od1.1 \*\*\**<br>*Od1.1 vs Od1.4 \*\*\**<br>*Od1.4 vs Od2 \*\*\** | F(5,120)= 0.94, p=0.46 |

Figure 6B shows the mean instantaneous respiratory frequency during the different periods of the first Post-Surgery session for both groups of rats: before the first presentation of odor 1 (Pre-Od1) and following each presentation of odor 1 (Od1.1, Od1.2, Od1.3 and Od1.4) and odor 2 (Od2). A two-factor ANOVA revealed that there was no effect of group nor interaction between group and period. However, there was a significant effect of the period (Table 4). Specifically, post-hoc analysis revealed a significant difference between Pre-Od1 and Od1.1 highlighting that both groups of rats detected Od1 with a similar sniffing frequency (8.27 and 8.36 Hz for 6-OHDA and Sham rats, respectively; Fig. 6B). In addition, we observed a significant decrease in the mean instantaneous sniffing frequency across the four Od1 presentations from 8.3Hz to 4.1Hz on average on both groups, indicating habituation to this odor. Finally, post-hoc comparisons revealed a significant difference between the last presentation of Od1 (Od1.4) and the presentation of the new odor (Od2) with a significant increase in the mean instantaneous respiration frequency (from 4.1Hz to 7.5 Hz), indicating that animals have detected, memorized and discriminated Od2 and Od1.

To summarize, across all sessions, 6-OHDA rats displayed similar odor-evoked sniffing in response to a new odor and to its repeated presentation compared to Sham rats. Therefore, the 6-OHDA treatment does not significantly alter high-frequency sniffing whether it is exploratory or odor-evoked.

## Discussion

The main aim of the present study was to test the potential impact of behavioral state on respiration deficits in a rat model of PD. To do so, we recorded and analyzed respiration under different behavioral states: during passive exposure to the experimental cage, leading to different vigilance states (sleep, quiet waking and exploration) and during an odor task assessing olfactory abilities. We first observed that 6-OHDA and Sham rats spent the same amount of time in the different vigilance states. Second, we tested whether respiratory alterations observed in rat model of PD could be specific of a given vigilance state. We did not observe any significant differences in respiratory parameters (instantaneous respiratory frequency and total amplitude) between 6-OHDA and Sham rats during sleep. However, we observed significant differences in respiratory parameters between 6-OHDA and Sham rats during wake (exploration and quiet waking). Finally, when sniffing behavior was elicited by odors in the odor task, we did not observe differences of respiratory frequency and olfactory capacities between the two groups.

It has been shown that respiratory patterns are different depending on the vigilance states and emotional content (Bagur et al. 2018, 2021; Dupin et al. 2020; Girin et al. 2021; Hegoburu et al. 2011). In addition, Janke et al. (2022) have recently demonstrated, using machine learning model, that behaviors in rodents can be accurately predicted based on their respiration patterns, therefore unraveling the strong correlation between the two parameters. In our study, we first verified whether the neurotoxin-induced lesion had altered the rats’ behavioral states. We showed that there was no difference in the total amount spent in each vigilance state (i.e. sleep, quiet waking and exploration) between 6-OHDA and Sham rats. Casarrubea et al. (2019) performed an extensive examination of ten different spontaneous behaviors and their sequence in an open-field in unilaterally lesioned 6-OHDA rats (21 days after the lesion). Comparatively to our findings, they have not observed any difference in the mean duration of each behavior (e.g. walking, shaking, immobility, climbing…). While there was no difference in the average duration of those behaviors, the authors still noted a decrease in the probability of occurrence of walking, immobile sniffing and stretched sniffing. Another interesting result from this study is the fact that 6-OHDA lesioned rats display a deficiency in their sequential behavioral organization: with less, shorter and less variable sequences of behaviors in 6-OHDA lesioned rats. It would be interesting in the future to associate a non-invasive respiration recording to the observation of this wide range of more complex behaviors expressed in the open-field.

### Effect of 6-OHDA treatment on respiration during different behavioral states

Given the highly specific association between respiratory patterns and behaviors, we tested the dissimilarities in respiratory parameters between 6-OHDA and Sham rats across the same behavioral state.

We did not observe significant differences in respiration during sleep between 6-OHDA and Sham rats. Our results are in accordance with studies showing no effect on respiration between 6-OHDA and Sham rats at rest (Andrzejewski et al. 2016, 2019). In these studies, no difference in respiratory frequency during normocapnic breathing was observed 14 days post unilateral medial forebrain bundle injection of 6-OHDA. Interestingly, rats were recorded when breathing around 100 breaths/min, corresponding to an approximate respiratory frequency of 1.7Hz. Our results have shown that this respiratory frequency is associated to a sleep state, therefore corroborating our findings.

Although we did not note any differences in respiratory parameters during sleep, we observed a higher respiratory frequency during quiet waking in 6-OHDA rats as well as a higher respiratory total amplitude during exploration and quiet waking. A previous study also showed an increase in baseline respiratory frequency in a rat model of PD. Indeed, Johnson et al. (2020b) used the Pink1−/− rat model of early-onset PD and showed higher respiratory frequency (approximately 2.25 Hz in 6 to 10-month-old Pink1-/-rats) compared to Sham rats (approximately 1.95 Hz) but no difference between groups on peak inspiratory or expiratory flows. They also mentioned in their methods that baseline was recorded once the rats were quiet but awake, which could be comparable to our definition of quiet waking state. However, there are other studies that showed that under normoxic conditions, 6-OHDA rats display a reduced respiratory frequency and minute ventilation compared to controls (Andrzejewski et al. 2020; Bialkowska et al. 2016; Falquetto et al. 2020; Fernandes-Junior et al. 2018; Lima et al. 2018; Oliveira et al. 2017; Tuppy et al. 2015). As highlighted by Andrzejewski et al. (2019), different experimental conditions as well as different PD models could explain those discrepancies: the site of injection of 6-OHDA (SNc, Medial Forebrain bundle or striatum), unilateral or bilateral lesions, the dose of 6-OHDA used (for example Tuppy et al. (2015) observed a reduced respiratory frequency in 6-OHDA rats only when using 12 or 24µg 6-OHDA in striatum but not for 6µg), the use of desipramine protecting noradrenergic and serotoninergic neurons from the neurotoxic effect of 6-OHDA, the timeline of the experiments (14, 21, 30, 40- or 60-days post lesion). In addition, the respiratory frequencies observed in those different studies in rats (Falquetto et al. 2020; Fernandes-Junior et al. 2018; Lima et al. 2018; Oliveira et al. 2017; Tuppy et al. 2015) are close to 100 breaths/min (∼1.7Hz) in Sham rats while values are even closed to 75 breaths/min (∼1.25 Hz) for 6-OHDA rats. Those values are similar to the ones we observed during sleep, suggesting that it is also possible that this difference in respiration frequency is ascribable to different behaviors (quiet/falling asleep vs. sleep) or a different stage of sleep in the two groups (Girin et al. 2021). This result highlights the need for future respiration-focused studies to include the analysis of behavior to circumvent a potential behavior bias between Sham and rodent model of PD.

6-OHDA treatment had an impact on respiratory cycle total amplitude during both exploration and quiet waking but the effect on respiration frequency was only observed during the quiet waking state. A first hypothesis is that this vigilance state difference in respiration could originate in differences of neuromodulator’s levels. Indeed, it has been shown that the response to CO_2_ is state-dependent due to the contribution of orexinergic and serotoninergic neurons (Nattie and Li 2012). Changes in these neuromodulators’ levels could therefore be responsible for the difference in respiration in a state dependent manner. Another possible explanation for the higher respiratory frequency observed in our 6-OHDA rats compared to Sham rats during quiet waking as well as higher total amplitude during both exploration and quiet waking can be related to anxiety. Indeed, 6-OHDA rats have been shown to display higher anxiety-related behaviors (Carnicella et al. 2014; Castrioto et al. 2016; Tadaiesky et al. 2008). Anxiety has been shown to modulate respiratory parameters (Boiten et al. 1994; Carnevali et al. 2013, 2014; Masaoka et al. 2003). Notably, Carnevali et al. (2013, 2014) have demonstrated that high-anxiety behavior rats display higher respiratory frequency than low-anxiety behavior rats. In addition, anxious rats display increased arousal (McAuley et al. 2009). Therefore, it cannot be excluded that during quiet waking, 6-OHDA rats are more anxious and/or display increased arousal and thus these behavioral features could contribute, at least to some extent, to a higher respiratory frequency.

Interestingly and relative to the effect on respiration frequency during quiet waking, Cavelli et al. (2019) recorded local field potentials in the olfactory bulb and primary motor cortex in a rat model of PD. The activity of the olfactory bulb, and notably gamma oscillations, is strongly shaped by respiration (Buonviso et al. 2006). The authors observed a decrease in gamma coherence between the olfactory bulb and primary motor cortex as well as changes in the respiratory-gamma coupling only during quiet waking but not during sleep. This effect could relate to changes in the respiratory parameters in 6-OHDA rats during this specific state of quiet waking.

### Odor-evoked respiratory responses and olfactory capacities in 6-OHDA rats

In a separate session, we used an odor paradigm allowing to trigger strong sniffing responses and to assess different olfactory functions. We showed that 6-OHDA rats display similar odor-evoked sniffing frequency in response to a new odor and its repeated presentation compared to Sham rats. This result means that sniffing frequency whether it is exploratory or triggered by an odor is not affected by the 6-OHDA treatment.

Sniffing has been demonstrated to be a reliable index of olfactory functions (Boulanger-Bertolus et al. 2023; Coronas-Samano et al. 2016; Courtiol et al. 2014; Dupin et al. 2020; Johnson et al. 2020a; Kepecs et al. 2006; Lefèvre et al. 2016; Rojas-Líbano and Kay 2012; Wesson et al. 2008, 2009; Youngentob et al. 1987). In the odor habituation/cross-habituation task, an increase in sniffing frequency between Pre-Od1 and Od1.1 is an index of odor detection, decrease in sniffing frequency across repeated presentations of a same odor is an index of odor habituation and increased sniffing frequency between the new odor and the fourth presentation of the first odor is an index of discrimination and of short-term memory processes (Al Koborssy et al. 2019; Coronas-Samano et al. 2016). Importantly, recording odor-evoked sniffing allowed us to investigate these four aspects without any bias related to possible motor deficits in 6-OHDA rats (Johnson et al. 2020a). In our experimental conditions, we observed, using this sniffing index, that 6-OHDA rats have similar olfactory capacities and short-term memory processes as Sham rats. In accordance with the present results and using the same PD model as ours, Drui et al. (2014) have shown that 6-OHDA rats do not display any olfactory deficit measured with attraction to an appetitive odor and avoidance to an aversive odor. In addition, Kurtenbach et al. (2013), using three different genetic mouse models of PD as well as mice treated with MPTP, have shown that those different mice models of PD have no major olfactory impairments. In addition, the different groups display similar electro-olfactogram responses to odors compared to wild-type mice. Other studies have reported some alterations of the olfactory capacities in rodent models of PD (Bonito-Oliva et al. 2014; Fleming et al. 2008; Ilkiw et al. 2019; Johnson et al. 2020a; Zhang et al. 2019). For example, Zhang et al. (2019) demonstrated that mice injected with 6-OHDA in the SN display reduced odor spatial memory and odor discrimination 7 days post-lesion but no alteration in odor detection nor habituation in an habituation/cross-habituation paradigm. Interestingly, Bonito-Oliva et al. (2014) demonstrated that mice with bilateral injection of 6-OHDA within the striatum show similar odor habituation as controls. However, they present reduced response to the new odor stimuli only when the previous odor belonged to the same category; those mice display preserved discrimination when the two odors were from different categories (e.g. non-social vs. social odor). Finally, using sniffing as an index of olfactory ability, Johnson et al. (2020a) demonstrated that female mice, but not male, model of PD display an altered odor detection sensitivity. Altogether, these studies reveal that in some experimental conditions, some olfactory deficits (notably discrimination or odor spatial memory) can be observed. The absence of effect on olfactory abilities assessed through the lens of sniffing might have been different if we had used either perceptually similar odors within the pair or/and lower concentrations or/and testing females. This result also highlights that olfaction is a really predominant sense for rodents and there is a need to challenge them with more complex tasks recruiting different olfactory skills and behaviors combined with respiration recording (e.g. telemetric jackets).

### Conclusion

To conclude, our results highlight some alterations of respiratory parameters during specific vigilance states in 6-OHDA rats notably during the awake states. This observation is important given the increasing number of studies demonstrating the importance of respiration for brain rhythms and cognitive functions. Indeed, respiration is believed to modulate neural oscillations of various brain regions (Heck et al. 2019; Juventin et al. 2023; Kluger and Gross 2021; Tort et al. 2018). Many of these cerebral structures are involved in diverse brain processes, such as cognition, stimulus processing, and emotion, whose expression partly depends on synchronous neuronal activity. The fact that we observed an effect of the SNc lesion on respiratory pattern during the quiet waking state is interesting since the large-scale synchronization of a majority of brain structures to the respiratory rhythm was mostly observed under a quiet waking state associated with low respiratory frequency (Girin et al. 2021). Therefore, disturbed breathing under a quiet state could interfere with a proper brainwave synchronization and result in associated cognitive deficits. Future studies will be needed and will have to combine brain rhythms recordings, respiration recordings and fine behavioral analysis.

## Acknowledgments

We would like to thank Ounsa Jelassi and Alexandre Valenti for their technical support on the project as well as Hayet Kouchi for helpful comments on the manuscript. We are very grateful to Sébastien Carnicella and Sabrina Boulet for their help in setting up the rodent model of PD in our lab.

